# Super Resolution Microscopy and Deep Learning Identify Zika Virus Reorganization of the Endoplasmic Reticulum

**DOI:** 10.1101/2020.05.12.091611

**Authors:** Rory K. M. Long, Kathleen P. Moriarty, Ben Cardoen, Guang Gao, A. Wayne Vogl, François Jean, Ghassan Hamarneh, Ivan R. Nabi

**Affiliations:** Life Sciences Institute, University of British Columbia, Vancouver, BC, Canada; Department of Cellular and Physiological Sciences, University of British Columbia, Vancouver, BC, Canada; School of Computing Science, Simon Fraser University, Burnaby, BC, Canada; Department of Microbiology and Immunology, University of British Columbia, Vancouver, BC, Canada; 5chool of Biomedical Engineering, University of British Columbia, Vancouver, BC, Canada

## Abstract

The endoplasmic reticulum (ER) is a complex subcellular organelle composed of diverse structures such as tubules, sheets and tubular matrices. Flaviviruses such as Zika virus (ZIKV) induce reorganization of endoplasmic reticulum (ER) membranes to facilitate viral replication. Here, using 3D super resolution microscopy, ZIKV infection is shown to induce the formation of dense tubular matrices associated with viral replication in the central ER. Viral non-structural proteins NS4B and NS2B associate with replication complexes within the ZIKV-induced tubular matrix and exhibit distinct ER distributions outside this central ER region. Deep neural networks trained to identify ZIKV-infected versus mock-infected cells successfully identified ZIKV-induced central ER tubular matrices as a determinant of viral infection. Super resolution microscopy and deep learning are therefore able to identify and localize morphological features of the ER and may be of use to screen for inhibitors of infection by ER-reorganizing viruses.

## Introduction

The endoplasmic reticulum (ER) is a highly dynamic network composed of 30-100 nm ribosome-studded rough ER sheets and convoluted networks of smooth ER tubules (1, 2). ER shaping proteins such as the lumenal sheet spacer protein cytoskeleton-linking membrane protein 63 (CLIMP-63), membrane curvature stabilizing reticulons (RTN) and junction stabilizing atlastin (ATL) are responsible for the generation of the morphologically distinct domains that comprise the ER (2–4). Recently, 2D STimulated Emission Depletion microscopy (STED) super-resolution microscopy combined with high speed live cell or rapid fixation approaches showed that RTN and CLIMP-63 regulate not only the ER macrostructure of rough ER sheets vs smooth ER tubules but also the nanodomain organization of sheets and tubules imaged in the cell periphery (5, 6). Increasing both 3D spatial and temporal resolution with various super resolution imaging techniques, including 3D structured illumination (SIM), grazing incidence structured illumination (GI-SIM) and lattice light sheet point accumulation for imaging in nanoscale topography (LLS-PAINT), showed that peripheral sheets were actually dense tubular matrices (7). While application of super-resolution microscopy to define morphology of peripheral ER (PER) structures has proven fruitful, its use to study the central ER (CER), and changes associated with cellular stressors has proven more challenging.

The ER is often utilized by pathogens to subvert the cytoplasmic innate immune response and promote replication (8–10). Bacteria, such as *Legionella pneumophila* and *Brucella abortus*, RNA viruses, and some intracellular protist pathogens, such as *Toxoplasma gondii*, all utilize the unique functions of the ER to promote their own replication (11–13). Flaviviruses, and coronaviruses, reorganize ER membranes into novel structures to better facilitate viral replication (12, 14, 15). A well-studied ER reorganizing flavivirus is the Zika virus (ZIKV), a member of the flavivirus genera within the *Flaviviridae* viral family that includes the dengue virus (DENV), and has shown causative association with congenital microcephaly and other neurological diseases (16–18). ZIKV-induced ER structures include replication factories, 60-100 nm spherical vesicular invaginations within the rough ER lumen that contain viral double-stranded RNA (dsRNA), an intermediate in genomic replication, that are the site of viral RNA genomic replication (19, 20). Associated with replication factories are ZIKV-induced convoluted networks of ER tubules, known as convoluted membranes, that are for the most part smooth although rough convoluted membranes containing replication factories have been reported in neural progenitor cells (19, 21).

The ZIKV RNA genome encodes 10 proteins, corresponding to the DENV genome, including 3 structural proteins (Capsid (C), Membrane (prM/M), and Envelope (E) proteins), involved in assembly of the virus capsid and envelope, and 7 non-structural (NS) proteins (NS1, NS2A, NS2B, NS3, NS4A, NS4B, and NS5) (22, 23). The NS proteins form the replication complexes and play key roles in the evasion of the host innate immune response, the hijacking of host cellular pathways, and the reorganization of ER membranes to promote viral replication (24). All of the flaviviral NS proteins are believed to play a role in the formation of the genomic replication machinery and replication factories. ZIKV NS3 and NS4B have been shown to overlap with convoluted membranes and replication factories by light microscopy (19, 25). NS2B is an integral membrane protein which localizes NS3 to replication factories and functions as a cofactor for the protease activity of NS3; NS2B and NS3 localize to both convoluted membranes and replication factories by electron microscopy (EM) (19, 26–28). NS4B is a transmembrane protein with five integral transmembrane domains, previously shown to be involved in the formation of convoluted membranes in DENV-infected Huh7 cells (26).

Virus-induced ER structures have been well-characterized by EM, however the cellular distribution of these structures and associated viral proteins remains poorly understood. Here, we use 3D STED super-resolution microscopy to show that ZIKV infection induces the reorganization of the CER to form dense tubular matrices associated with ZIKV replication sites. We further apply deep learning approaches to identify ZIKV-infected cells based on changes in ER morphology. Quantitative detection of virus-induced changes to the ER represents proof-of-principle for the application of deep learning-based image analysis of ER reorganization for large scale drug screening and identification of compounds inhibitory to viral infection.

## Results

### ZIKV infection induces the formation of a dense tubular matrix in the CER

In order to study ER morphology following ZIKV infection, we first generated stable U87 glioblastoma cells transfected with either ER monomeric oxidizing environment-optimized green fluorescent protein (ERmoxGFP), a lumenal ER reporter containing the bovine prolactin signal sequence and KDEL ER retention sequence linked to inert, monomeric, cysteine-less moxGFP (29), or the membrane-associated ER reporter Sec61β-GFP. U87 cells stably expressing the ER reporters were then infected with ZIKV strain PRVABC59 (Puerto Rico 2015) at a multiplicity of infection (MOI) of 1 for 48 hours. Cells were fixed with 3% paraformaldehyde/0.2% glutaraldehyde to preserve ER architecture (6, 7, 30, 31) and labeled for dsRNA, a marker for ZIKV replication factories (19). Maximum projections of 3D STED image stacks show high intensity ERmoxGFP and Sec61β-GFP labeling in a CER region and low intensity labeling in PER tubules in mock-infected cells (Figure 1A), as reported previously by diffraction limited confocal microscopy (3). Upon ZIKV infection, the CER reorganizes to form an intensely labeled crescent-shaped region surrounding a lower intensity perinuclear region (Figure 1A). Interestingly, the crescent-shaped ZIKV-induced perinuclear ER overlapped extensively with dsRNA (Figure 1A). Imaris Bitplane software fragments the ER into distinct segments that can then be analyzed for different features, including reporter density, segment Z-height and segment overlap with other labels, such as dsRNA. Density-based segmentation of the ERmoxGFP- and Sec61β-GFP-labelled ER of ZIKV-infected cells showed that the higher density crescent-shaped CER region exhibited significant overlap with dsRNA-positive ER structures relative to the rest of the ER (Figure 1B). This suggests that ZIKV dsRNA associates with an ER region of high density for both lumenal and membrane ER reporters.

**Figure 1.**
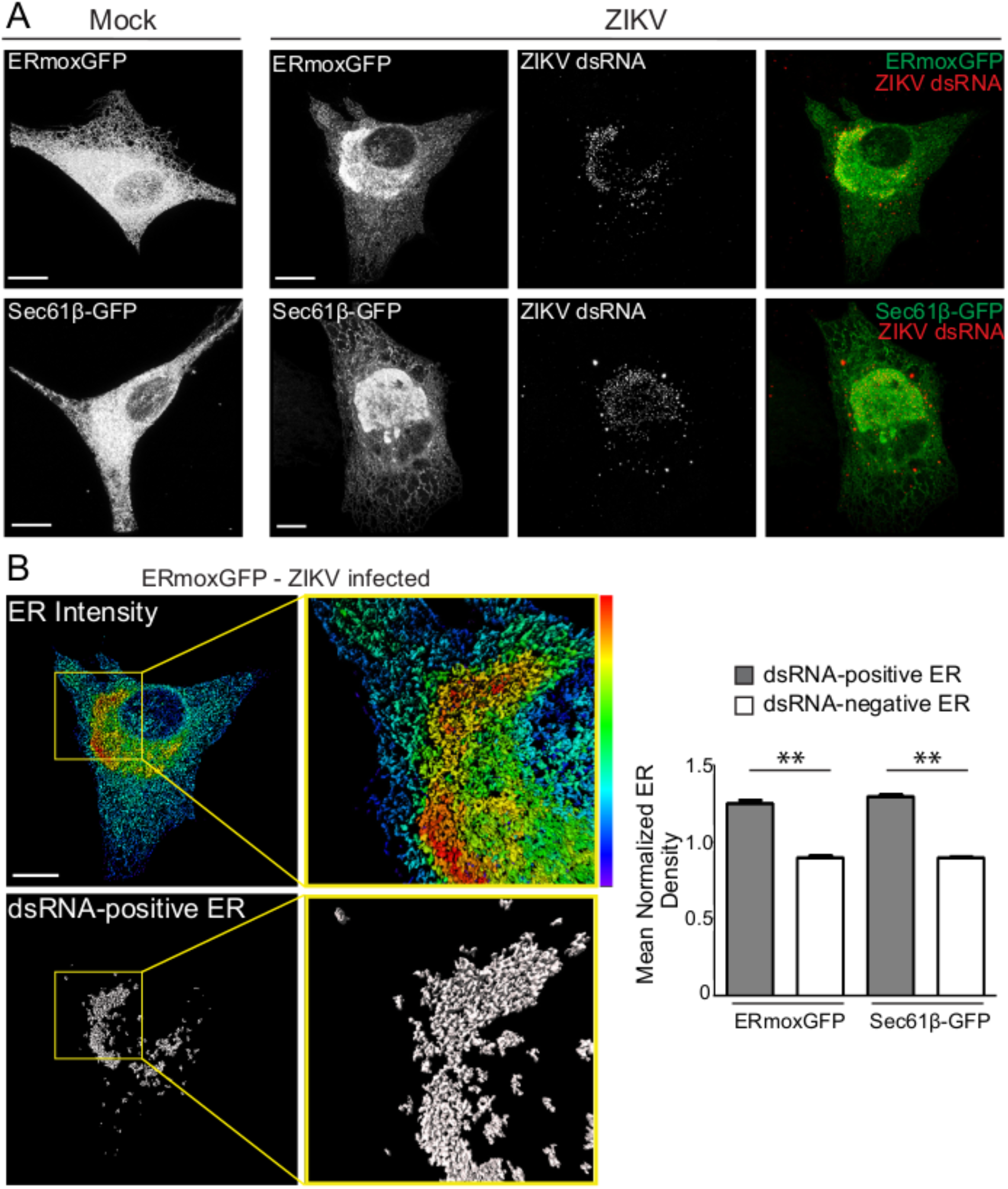
3D STED microscopy reveals ZIKV-induced ER reorganization in human U87 glioblastoma cells. A) ERmoxGFP or Sec61β-GFP stably transfected U87 cells were mock-infected or infected for 48 hours with the PRVABC59 2015 ZIKV strain (MOI= 1). ER reporter GFP and immunostained dsRNA-labeled ZIKV replication factories were imaged by 3D STED microscopy. B) Fluorescence intensity of ERmoxGFP of infected cells using a spectrum heat map and a segmentation mask of the ER that colocalizes with dsRNA (grey), both generated on Imaris x64 9.2.1 (Imaris), are depicted. Yellow squares in the panels indicate the magnified ROIs shown in the adjacent panels. Quantification of the mean normalized ER density ((Intensity sum of mask/total cell intensity sum)/ (volume sum of mask/ total cell volume sum)) was performed for both dsRNA-positive and dsRNA-negative ERmoxGFP and Sec61β-GFP in ZIKV-infected cells by Imaris segmentation. Scale bar= 10 microns. 5 cells per biological replicate (N=3). Statistics were done using unpaired Student’s T tests: **= P<0.01, Error bars represent SEM.

We then investigated the relationship between the dense ZIKV-induced crescent-like region and the CER and PER. Segmenting the ER based on Z-height of ER segments showed that PER tubules were present in 3-5 consecutive frames of 210 nm each while CER regions were abundantly present across 7-10 consecutive frames (Figure 2A). For both ERmoxGFP and Sec61β-GFP labeled cells, average max Z-height of PER tubular regions was 0.95 μm and for CER regions 1.8 μm (Figure 2A). No PER segments reached a height greater than 1.26 μm. ER segments with a height above 1.26 μm were therefore classified as CER and ER segments with a height below 1.26 μm as PER, effectively segmenting the ER into CER and PER regions (Figure 2B). Based on this ER segment height-based classification of CER and PER, the CER was found to present a two-fold increase in volume and increased density relative to the PER (Figure 2C). ER density in the height based CER was elevated relative to PER and ZIKV infection showed minimal impact on the relative volume or density of CER and PER regions (Figure 2D).

**Figure 2.**
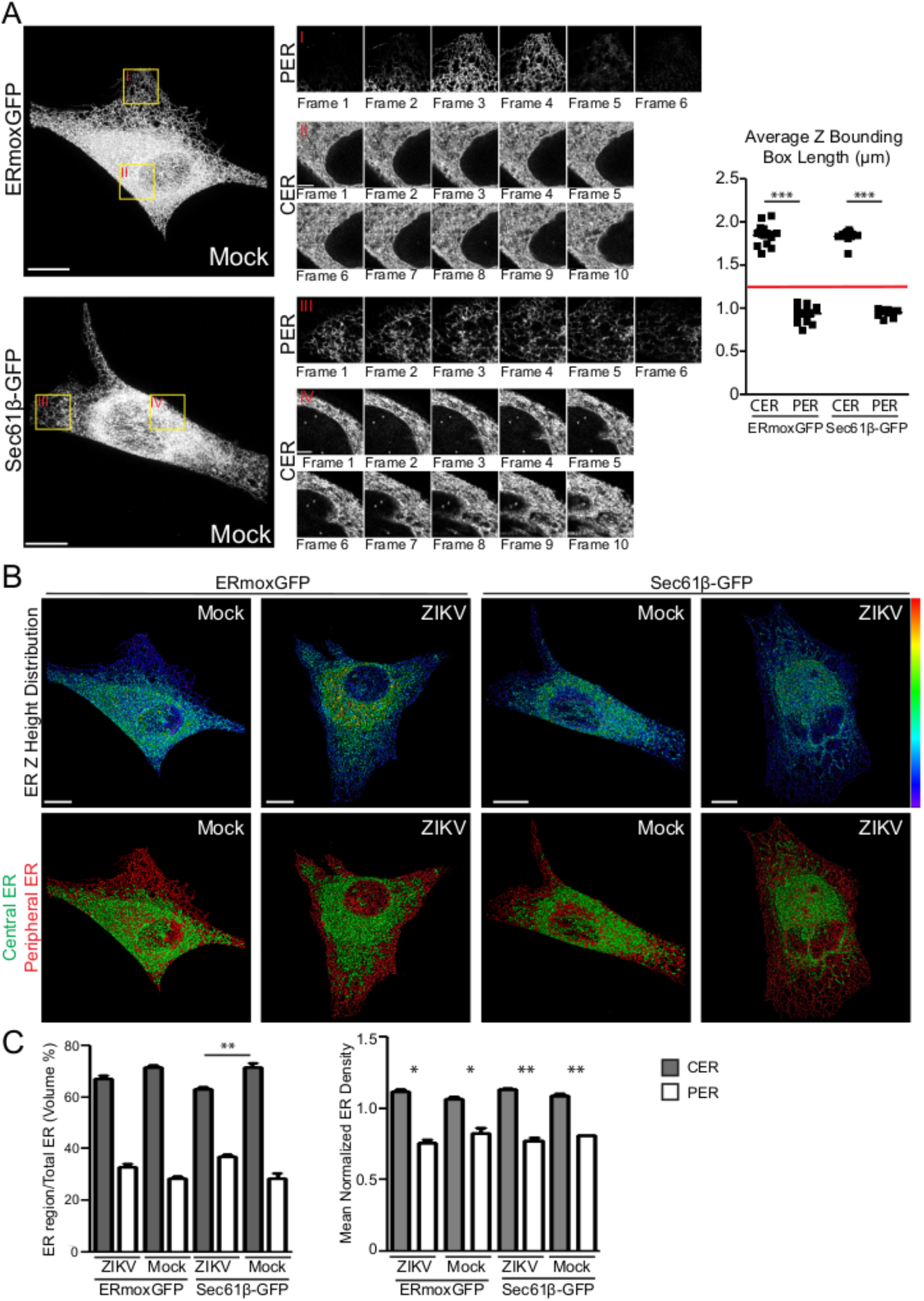
Defining the CER and PER of ZIKV- and mock-infected cells. A) Mock-infected ERmoxGFP and Sec61β-GFP stably transfected U87 cells were imaged by 3D STED microscopy. Magnified ROIs (yellow ROIs identified by red Roman numerals) show that the PER extends over 3-5 sections (210 nm step size) and CER >10 sections. Graph shows average PER and CER Z-height for each ERmoxGFP or Sec61β-GFP labeled cell. A Z-height cutoff of 1.26 microns (red line) was used to identify PER and CER objects. B) Segmented ER labeling from 48-hour ZIKV- or mock-infected ERmoxGFP or Sec61β-GFP stably transfected U87 cells (MOI= 1) was visualized using a Z-height spectrum heat map and CER (green; > 1.26 μm) and PER (red; < 1.26 μm) masks are shown. C) Volume percentage (left) and mean normalized density (right) of CER and PER masks between mock- and ZIKV-infected cells. 5 cells per biological replicate were analyzed for a total of N=3. ANOVA with post-hoc Tukey HSD: *= P<0.05, **= P<0.01, and ***= P*<0.001. Error bars represent SEM. Scale bar= 10 microns.

Overlaying the CER and PER masks with the dsRNA-positive ER mask showed that the dsRNA-positive ER (>80% volume/volume) is predominantly included within the CER mask (Figure 3A). Indeed, only 10% of PER volume contains dsRNA while 35% of CER volume contains dsRNA for both ER reporters (Figure 3A). Morphological comparison of the dsRNA-positive and -negative CER of ZIKV-infected cells with the CER of mock-infected cells showed that the CER was composed of a convoluted network of tubules for both the ERmoxGFP- and Sec61β-labeled ER (Figure 3B). 3D reconstructions confirmed that these regions were predominantly tubular with a few small sheet-like structures, similar to the tubular matrix morphology of peripheral sheets (7). 3D voxel-based visualization and quantification showed that the density of ER tubular structures in the dsRNA-positive ER is higher, for both the ERmoxGFP or Sec61β-GFP ER reporters, than in the dsRNA-negative CER regions of ZIKV-infected cells or the CER of mock-infected cells (Figure 3B). The lower ER reporter density reflects reduced spacing between tubules in the dsRNA-positive ER, suggesting that ZIKV infection induces tubular matrix reorganization in a subdomain of the CER in U87 cells. Consistently, EM analysis of the microcephaly relevant cerebral brain organoid model (32) showed that ZIKV-induced ER reorganization from perinuclear stacked rough ER sheets to a perinuclear, circular region of convoluted smooth ER tubules (Figure 4). These results are consistent with ZIKV induction of a perinuclear tubular matrix.

**Figure 3.**
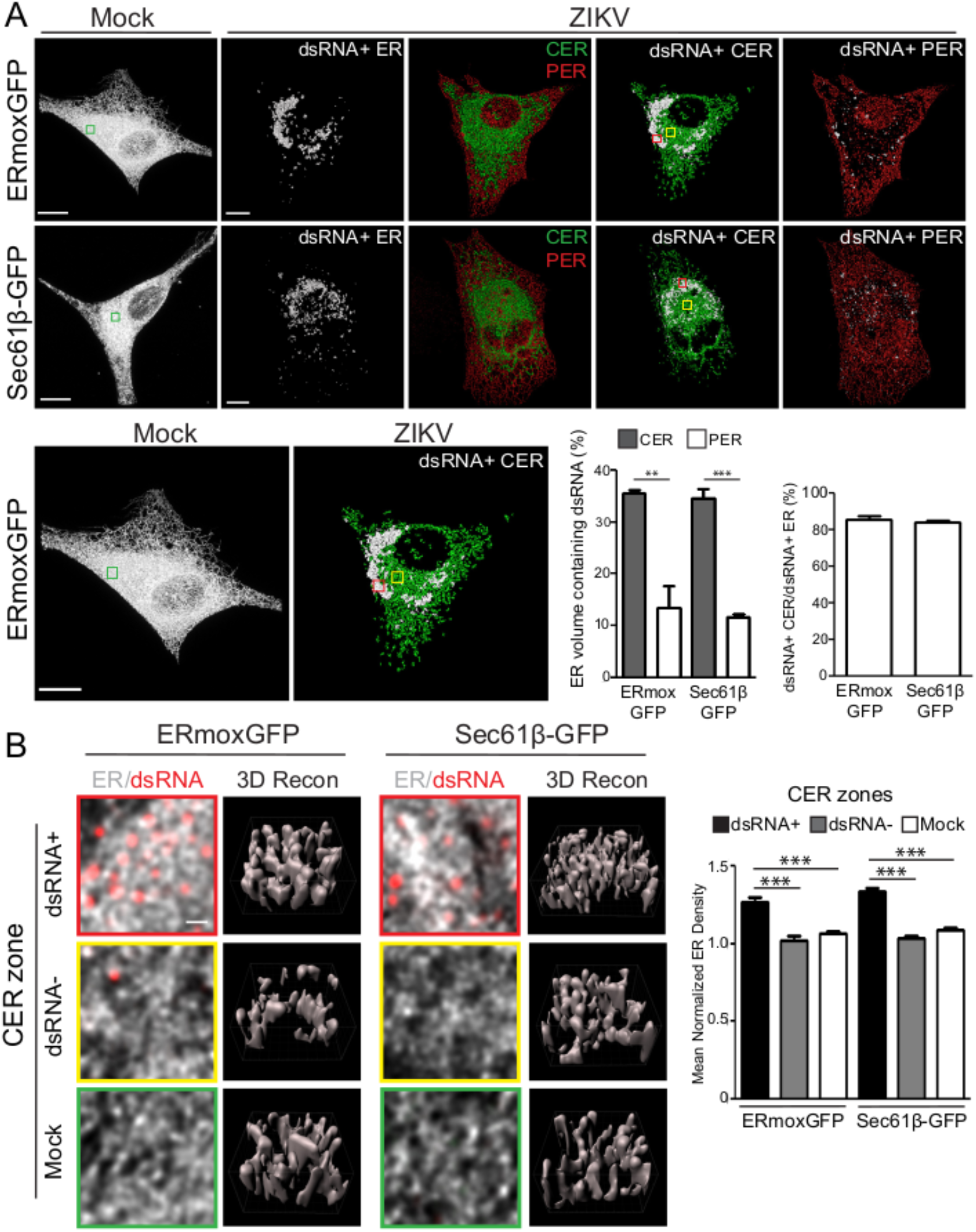
The ZIKV-induced dsRNA-positive CER region is a dense tubular matrix. A) Shown are representative mock-infected cells and CER (green) and PER (red) masks overlaid with the dsRNA-positive ER mask (white) for ZIKV-infected ERmoxGFP and Sec61β-GFP transfected U87 cells. Enlarged images of ERmoxGFP transfected cells show 3×3 μm ROIs of the mock-infected CER (green box) and of the ZIKV-infected dsRNA-positive (red box) and dsRNA-negative (yellow box) CER shown in B. Graphs show the volume percent of the CER or PER region that contains dsRNA-positive ER (left) and the volume percent of the dsRNA-positive ER that resides within the CER mask. B) 2D images of ER (white) and dsRNA (red) labeling in 3×3 μm ROIs of the ZIKV-infected dsRNA-positive (red) and dsRNA-negative (yellow) CER and mock-infected (green) CER are shown above Imaris 3D surface rendering of 1×1 μm regions of the above ROIs. Graph shows mean normalized ER density for each of the three CER zones by Imaris segmentation and masking. 5 cells per biological replicate were analyzed for a total of N=3. ANOVA with post-hoc Tukey HSD: *= P<0.05, **= P<0.01, and ***= P<0.001. Error bars represent SEM. Scale bar: 10 microns (500 nm for zoomed ROIs).

**Figure 4.**
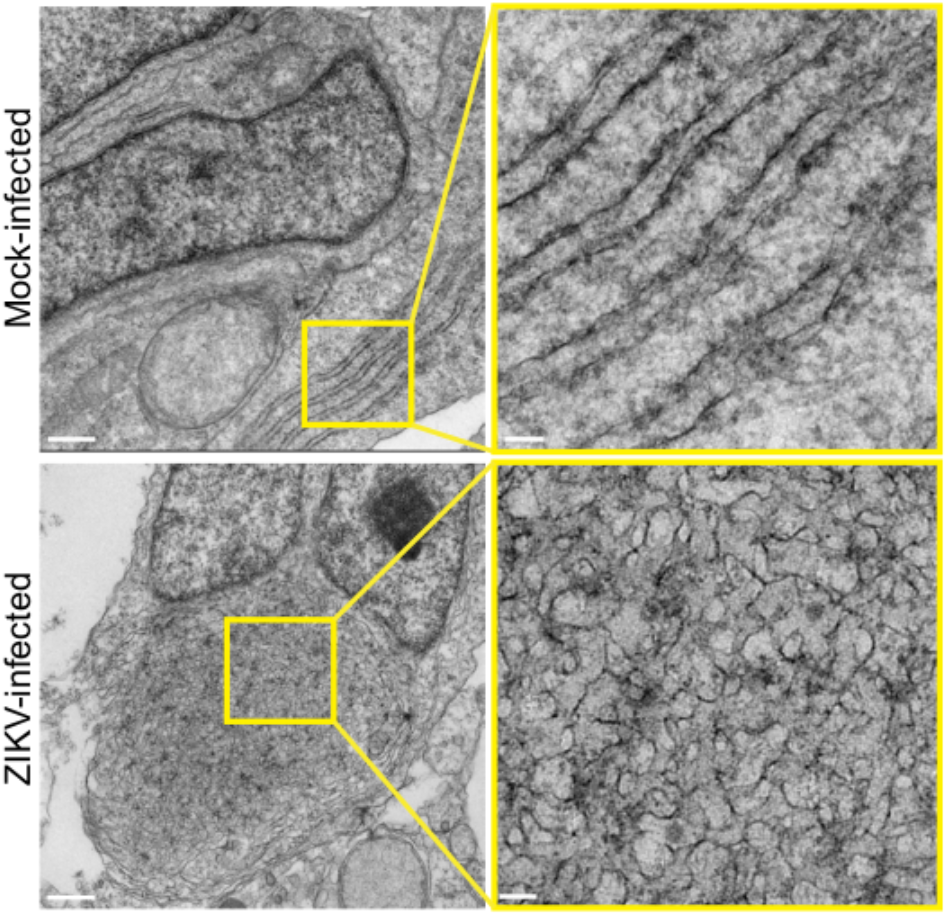
Ultrastructural analysis of ZIKV-infected cerebral brain organoids. Transmission EM images of 50 nm thin sections of 48-hour mock- and ZIKV-infected cerebral brain organoids (MOI=1). Yellow boxes show ROIs shown of adjacent higher magnification images that highlight rough ER sheets in mock-infected and tubular matrices (convoluted membranes) in ZIKV-infected cells. Scale bars: 500 nm and 100 nm for zoomed image ROIs.

### ER localization of ZIKV NS2B and NS4B structural proteins

We then labeled cells for ZIKV NS proteins NS2B and NS4B to assess their relationship to the ZIKV-induced tubular matrix. 3D STED analysis showed a predominant distribution of both NS2B and NS4B to the CER and more particularly to the dense ZIKV-induced crescent-shaped tubular matrix in ERmoxGFP transfected U87 cells (Figure 5A). While NS2B is predominantly associated with the dsRNA-positive CER, NS4B labeling extended throughout the CER as well as to the PER (Figure 5A). To quantify this, we identified NS2B-positive and NS4B-positive ER segments and determined their overlap with total ER, CER and PER (Figure 5A). While NS4B was present at high levels on both PER and CER segments, NS2B was enriched in the CER relative to the PER and presented a similar ER distribution to dsRNA (Figure 5A).

**Figure 5.**
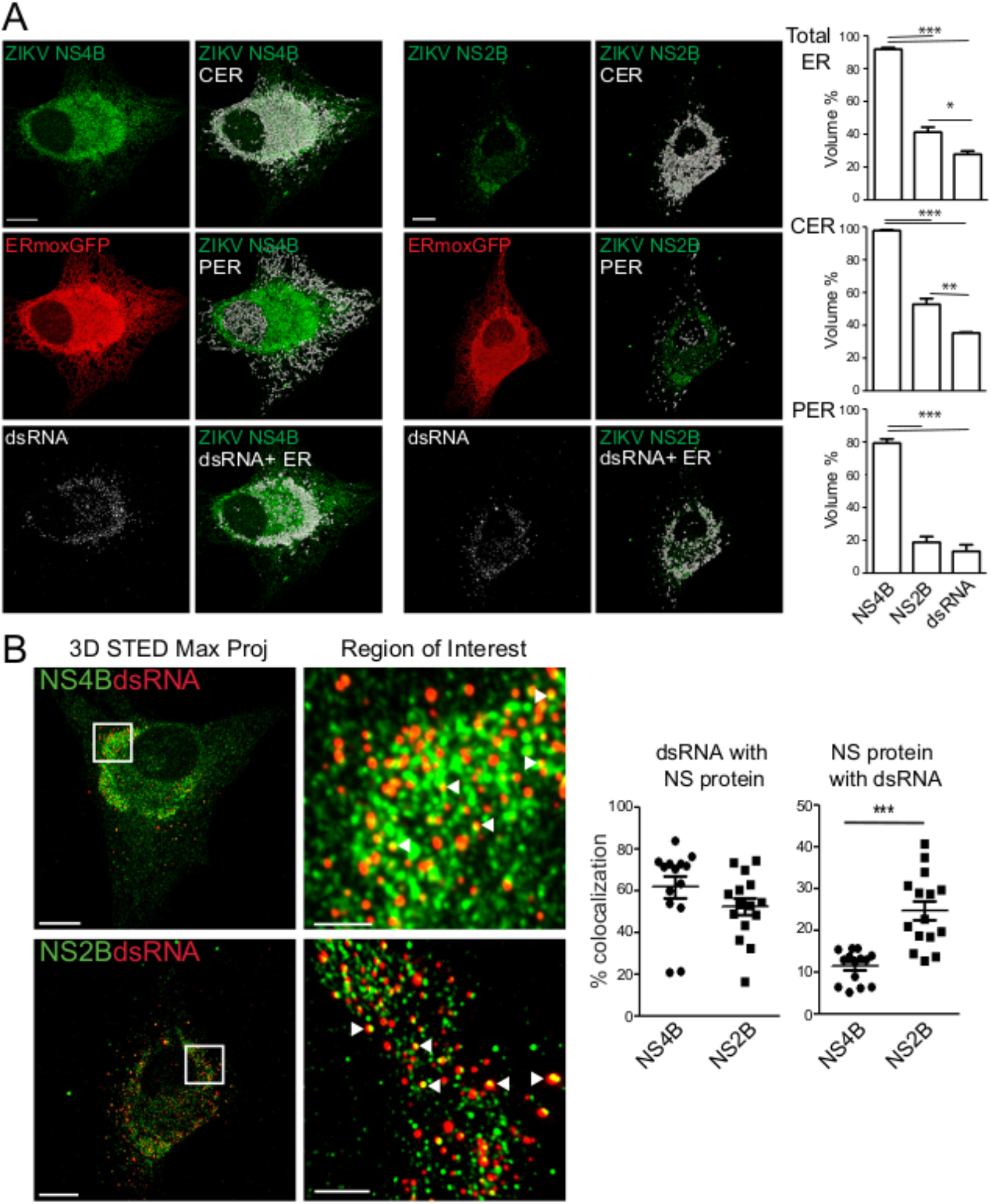
ER distribution of ZIKV NS2B and NS4B proteins. A) Mock- or ZIKV-infected ERmoxGFP (red) stably transfected U87 cells were immunostained for ZIKV NS2B or NS4B (green) and dsRNA (white). Merged images show NS2B or NS4B (green) overlaid with the CER, PER or dsRNA-positive ER (white). Graphs show quantification of volume percent of NS4B, NS2B and dsRNA ER regions relative to total ER, CER or PER. B) Mock- or ZIKV-infected ERmoxGFP stably transfected U87 cells were immunostained for ZIKV NS4B or NS2B (green) and dsRNA (red). White squares show ROIs of adjacent zoomed images in which white arrowheads show colocalization between NS protein and dsRNA puncta. Graphs show percent of dsRNA puncta overlapping NS4B or NS2B puncta (left) and percent of NS4B or NS2B puncta overlapping dsRNA puncta. 5 cells per biological replicate (N=3) with each dot representing a cell. ANOVA with post-hoc Tukey HSD: *=P<0.05, **=P<0.01, and ***=P<0.001. Error bars represent SEM. All images are maximum projections from 3D STED stacks. Scale bar= 10 microns. ROI scale bar = 2 microns.

The majority (>55%) of dsRNA-labeled puncta were associated with NS2B or NS4B, consistent with the presence of both these NS proteins in the ZIKV-induced tubular matrix. In contrast, a minority of NS2B (~25%) and NS4B (~10%) spots overlapped with dsRNA spots (Figure 5B). In the dsRNA-positive CER, the highly punctate NS2B labeling differed from a more reticular NS4B labeling. These two ZIKV NS proteins therefore exhibit distinct distributions within the ZIKV-induced tubular matrix when not associated with dsRNA replication complexes (Figure 5B). Together with the differential distribution of NS2B and NS4B within the ER as a whole (Figure 5A), these results highlight that these two ZIKV NS proteins do not associate exclusively with replication factories and suggest that following synthesis of the ZIKV polyprotein, NS2B and NS4B undergo distinct biosynthetic pathways before reuniting in ER-associated replication complexes.

### Deep learning identifies ZIKV-induced ER reorganization

Deep learning has been successfully applied to the task of image recognition, distinguishing the category or class to which a given image belongs. Deep learning architectures have outperformed shallow architectures when benchmarked on ImageNet a dataset widely used by the computer vision community that contains roughly 14 million images belonging to 20,000 classes (e.g., cats, dogs, plant species, various modes of transportation) (33–35). Deep convolutional neural networks (CNNs) are capable of learning local and global spatial patterns from raw training data (i.e. pairs of images and corresponding classes) enabling inference of the correct class of unseen images, and have achieved state of the art performance when benchmarked on ImageNet (36). Importantly, compared to non-deep learning detection methods such as machine learning, deep learning approaches: 1) avoid the need to design and select features for highly complex ER structures that are required by non-deep learning methods; and 2) provide the ability to move beyond simple classification to inspect discriminative regions (i.e. subregions of the ER within each cell).

We therefore applied deep neural networks to identify and distinguish the morphological features of the ER of ZIKV-infected cells. A pipeline outlining our approach is shown in Figure 6A. We train a CNN using 2D frames (each representing a single Z-frame) from 3D STED volumes of ERmoxGFP and Sec61*B*-GFP labeled ZIKV- and mock-infected cells. Our CNN builds off of VGG16, a deep neural network architecture proposed by Simonyan et al. (35) which achieved state of the art results on the 2014 ImageNet Large Scale Visual Recognition Challenge (ILSVRC) (37, 38). To improve computation time, we initially performed the analysis on downsampled STED images. As the 3D STED data sets were relatively small to train the CNN from scratch, we leveraged a network already pretrained on natural images. This transfer learning technique speeds up the convergence rate of networks when dealing with small target datasets. Certain filters (combinations of weights) learned on the first dataset (i.e. ImageNet) may still be useful for classifying a second dataset (i.e. STED); as a result, less weight updates are needed before achieving good performance. Using a pretrained VGG16 as our base model we obtained a 20% boost in test accuracy, compared against a random weight initialization. Using ERmoxGFP labelled ER alone, the CNN was able to distinguish between ZIKV- and mock-infected cells with an 82% accuracy (Figure 6B, top left). Accuracy using Sec61β-GFP labelled ER was only 66%, whereas combining cells labeled with both ER markers, increasing the number of training samples seen by the network, still resulted in only 71% accuracy. Based on the improved classification performance, further analysis was performed using ERmoxGFP labelled cells.

**Figure 6.**
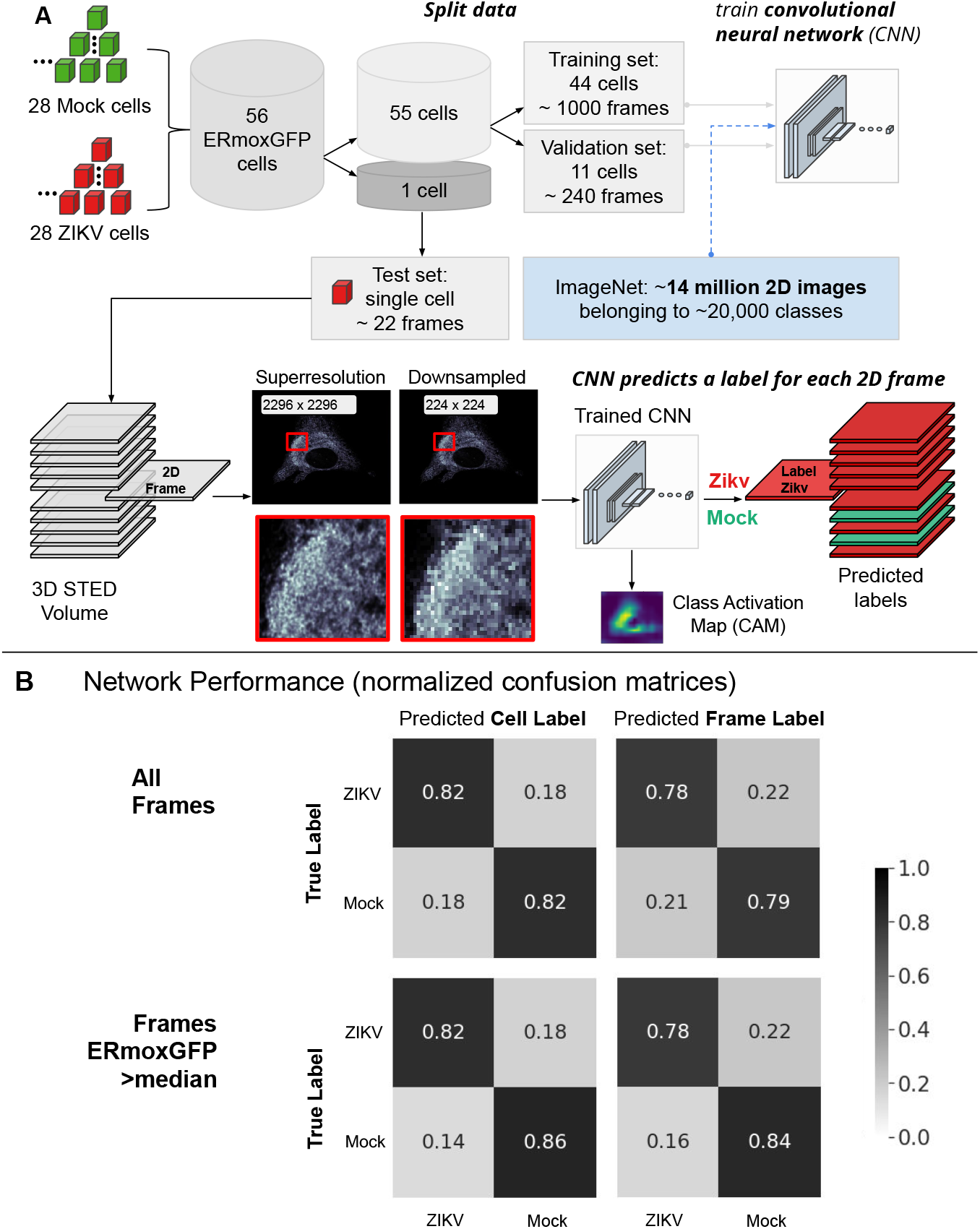
Deep learning classification pipeline: Pretrained convolutional neural network accurately predicts labels of 2D frames from 3D STED volumes. A) Leave-one-out cross validation is successively applied to each cell. This prevents information from 2D frames leaking between training, validation and test sets. During training, network uses 2D frames from 55 cells (specifically, 44 for training and 11 for validation). The trained CNN then predicts a class label (i.e. ZIKV-infected or mock-infected) to each 2D frame of the remaining test cell. Class Activation Maps are also generated for each 2D frame belonging to the test set. B) CNN performance reported on a cell basis and across 2D frames. Normalized confusion matrices report the total number of predicted labels (ZIKV-infected or mock-infected) over the total number of ground truth labels. For example, 79% of all mock-infected 2D frames were predicted correctly by the CNN (top right). Predicted cell labels correspond to the majority label of predicted frame labels for each cell (top left). When excluding frames beyond the cell with reduced ERmoxGFP signal, performance metrics increase both in terms of cell label predictions (bottom left) and individual frame label predictions (bottom right).

With a limited sample size of 56 ERmoxGFP labelled cells, we applied leave-one-out cross validation; 56 cells were split into a set of 55 cells used to train the CNN, and a test set consisting of the remaining cell. This process is outlined in Figure 6A, and is repeatedly applied using each cell as the test set. The 55 cells used to train the CNN are further split into a training and validation set, composed of 44 and 11 cells, respectively. During training, all 2D frames from cells belonging to the training set are passed individually to the CNN while 2D frames of the validation set cells are used to evaluate the performance and update the network’s parameters (weights) accordingly. This grouping of cells into training, validation and test sets ensures that information is not leaked, i.e., the CNN is only presented with unseen 2D frames during testing. ZIKV- and mock-infected class activation maps (CAMs) (39) are generated for each 2D frame to help define regions used by the CNN when inferring a class label (ZIKV or Mock). We achieved 82% accuracy and 82% specificity to identify ZIKV- and mock-infected cells based on a majority of frame predictions per cell. STED Z stacks extend beyond the cell and include Z sections that contain minimal ERmoxGFP signal and show poor accuracy to predict class label (Supp Fig 1). When considering those frames containing ERmoxGFP signal intensity greater than the median, we achieved 84% accuracy and 86% sensitivity. On a per frame basis, accuracy for all frames was 78% and sensitivity 79% that increased to 81% and 84%, respectively, when considering frames expressing ERmoxGFP greater than median intensity (Figure 6B, Supp. Fig. 1). Considering the confusion matrix for all frames, cell label (Figure 6B, top left), the 82% of ZIKV-infected 2D frames correctly predicted to be infected represent true positives (TP) while the 18% of ZIKV-infected 2D frames not predicted to be infected represent false negatives (FN). The 82% of mock-infected 2D frames correctly predicted to be uninfected represent true negatives (TN) and the 18% of mock-infected 2D frames predicted to be ZIKV-infected represent false positives (FP). The CNN has therefore accurately predicted which cells are ZIKV-infected based on the reorganization of ERmoxGFP-labeled ER. Improved accuracy for frames expressing higher ERmoxGFP highlights that the network is using the ERmoxGFP label to identify ZIKV-infected frames and cells.

To determine the basis for CNN decision making we analyzed the respective CAMs for both classes, ZIKV- and mock-infected cells. This method, first proposed by (39) produces a heatmap (or CAM), corresponding to a given image input, which can help localize the discriminating regions of the image that the CNN uses to infer a class label (Figure 7A). For instance, identification of ZIKV-infected slices (true positive; TP; orange) is primarily based on regions highlighted by the ZIKV CAM while identification of mock-infected cells (true negative; TN; green) is based on the mock CAM (Figure 7B). The range of values of the generated CAMs is between 0 and 1. Class activation maps have been used for weakly supervised segmentation, i.e. to delineate regions of interest (ROIs) (40, 41)We consider regions of interest (ROIs) to be areas of the CAM with values greater than 80% of the maximum. For example, if a CAM has maximum value of 1, the CAM thresholded at 80% considers only CAM regions with values greater than 0.8, corresponding to regions of high network attention. Representative examples of ZIKV (red) and mock (blue) CAMs are shown in Figure 7B; as they transition to yellow, concentric rings correspond to smaller CAMs and increasing thresholds from 10-95%. In general, the mock CAM was centered over the cell for correctly labelled uninfected cells (TN) cells as was the ZIKV CAM for correctly labeled ZIKV-infected cells (TP) cells (Figure 7B). Representative downsampled ERmoxGFP-labeled images from true positive (TP) instances show the 80% ZIKV CAM in red covering the central ER region. Higher resolution TP patches (112×112 pixels) from these regions of high network attention (ZIKV CAM with a threshold of 80%) show ER structures used to correctly label ZIKV-infected cells (Figure 7C).

**Figure 7.**
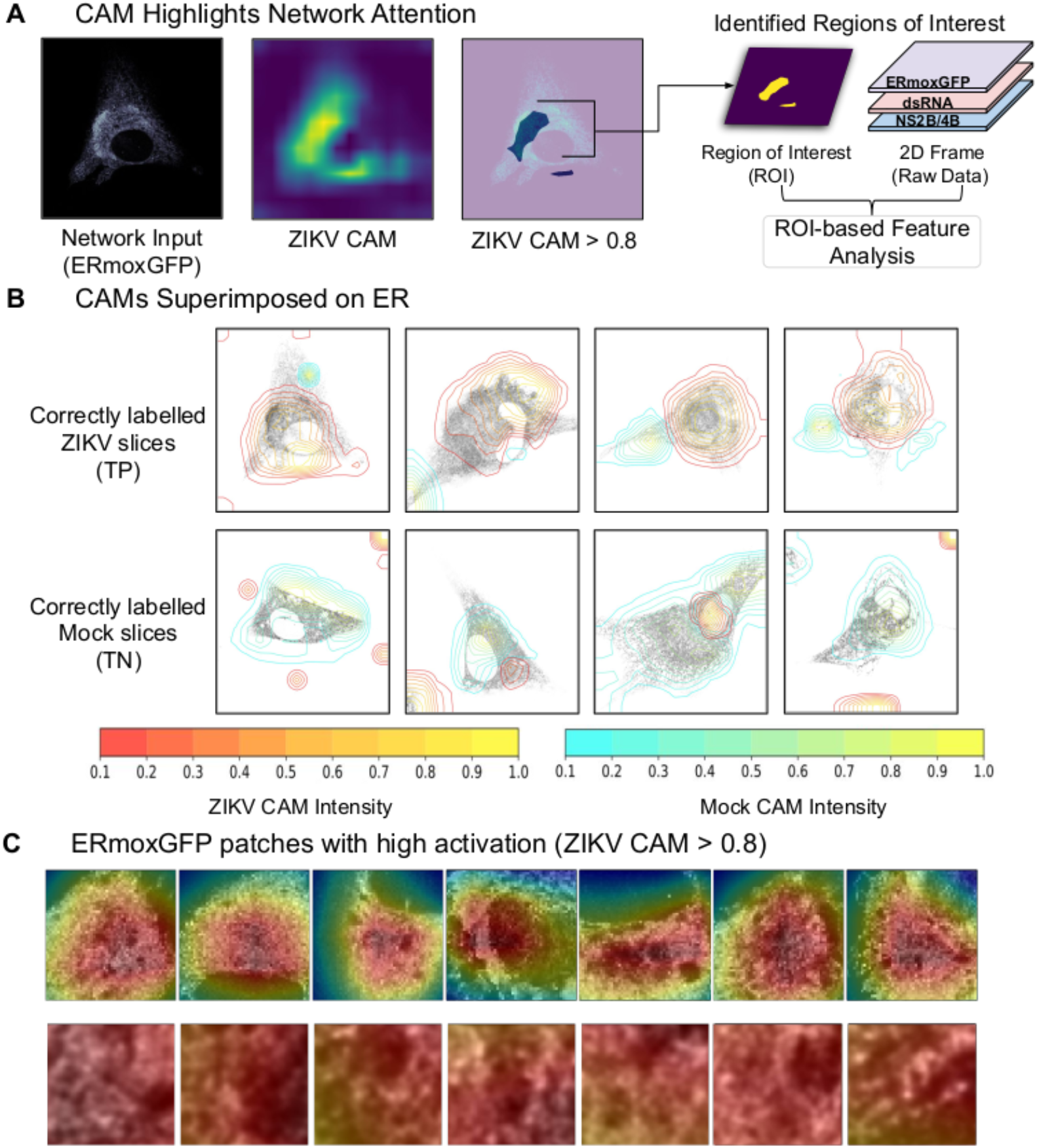
Class Activation Mapping for Region of Interest Based Feature Analysis. A) For each test 2D frame passed as input to the CNN, we generated Class Activation Maps (CAM) which are then thresholded to create ROIs. These regions identified by deep learning are then used for further analysis of ERmoxGFP, dsRNA and NS2B/4B. B) Contour maps of CAMs superimposed on ERmoxGFP labelled ER show regions identified by CAMs. The contour color represents varying CAM intensity from 0.1 (low network attention) to 1.0 (high network attention). For correctly labelled ZIKV-infected and mock-infected 2D frames, we show that the respective CAM focuses on the cell. C) The top row shows representative downsampled images from true positive (TP) instances highlighting the 0.8 ZIKV CAM in red. The bottom row shows 112×112 pixel patches from high resolution STED images within the 0.8 ZIKV CAM region.

### Deep learning identifies ZIKV-infected cells based on the high density CER

As seen in Figure 8A, ROIs identified by the ZIKV CAM show consistently higher ER density at all thresholds when comparing correctly identified infected frames (TP, orange) to the mock-infected frames (TN, green). Similarly, for 2D frames correctly identified as mock-infected (TN), the regions found by the mock CAM show increased ER density at all thresholds compared to the TP frames. This suggests that the CAMs used to identify both ZIKV- and mock-infected cells correspond to high ER density regions localized over the cell (see Figure 7B) and that VGG16 is using differences in the ER label (ERmoxGFP) to identify slices as either ZIKV- or mock-infected. Consistently, the density profile of the ZIKV CAM for cells falsely identified to be infected (false positive; FP; blue) matched that of TP (orange) ZIKV-infected cells while that of the mock CAM for cells falsely identified to be uninfected (false negative; FN; purple) matched the mock TN (green) profile. ER density for the ZIKV and mock TP and TN profiles at a CAM threshold of 80% show clearly that regions identified by the thresholded ZIKV CAM have significantly increased ER density when comparing TP to TN frames (Figure 8B, left). Conversely, regions identified by the thresholded mock CAM have increased ER density for TN compared to TP cells (Figure 8B, right). We then calculated the average Euclidean distance between the weighted center of mass of the CAMs and the weighted center of mass for all ERmoxGFP labelled pixels (Figure 8C). The center of mass of the ZIKV CAM for TP and for the mock CAM for TN were closest to the ERmoxGFP center of mass. As for the ER density profile of the CAMs, Euclidean distance of FPs for the ZIKV CAM matched TP, and FNs for the mock CAM matched TNs. Together, these data argue that the CAMs used to identify ZIKV- and mock-infected cells are located in dense ER regions near the center of mass of the ERmoxGFP label.

**Figure 8.**
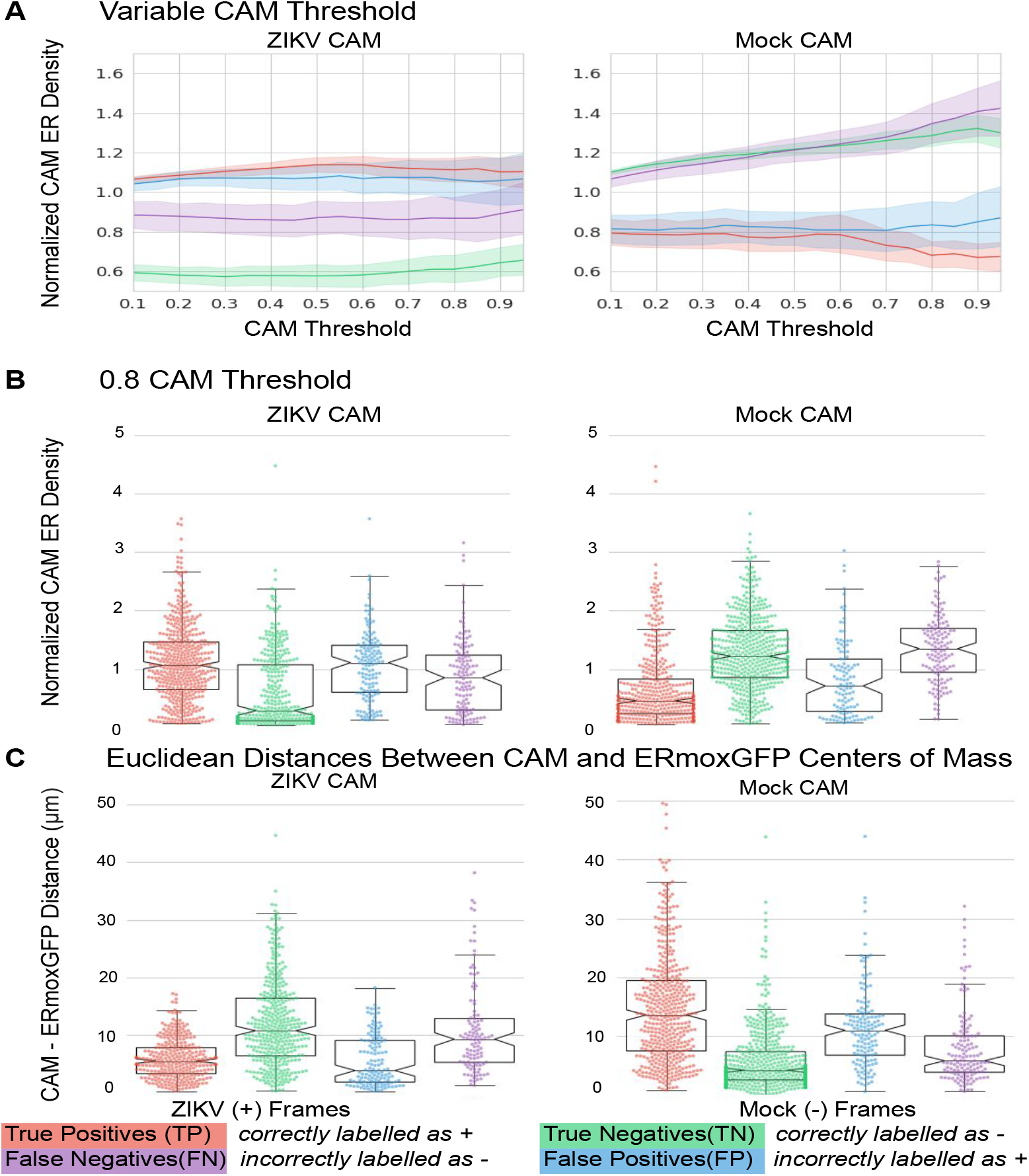
Normalized CAM ER density across subgroups. ROIs are defined by various CAM thresholds. For a given ROI, ER density is defined as total ERmoxGFP intensity within the ROI divided by ERMoxGFP area inside the ROI. ER density for each ROI defined by the CAM is then normalized by the ER density of the whole cell. A) ER density of ROIs defined by CAM thresholds from 10-95% with increments of 5% is compared across 4 subgroups: ZIKV-infected 2D frames correctly predicted to be infected (true positives); mock-infected 2D frames correctly predicted to be uninfected (true negatives); ZIKV-infected 2D frames incorrectly predicted to be uninfected (false negatives); mock-infected 2D frames incorrectly predicted to be infected (false positives). B) ER densities of 80% CAMs ROIs compared across subgroups. C) Euclidean distances between center of mass of 80% CAMs ROIs and weighted center of mass of ERmoxGFP signal.

To determine the relationship of the ZIKV CAM to the dense tubular matrix region that we identified by 3D STED super-resolution microscopy (Figure 3), we assessed CAM overlap with dsRNA, NS2B, NS4B and ERmoxGFP labeling. To do this, we calculated the normalized CAM intensity, defined as the intensity sum of a given channel in the ZIKV CAM normalized by the total area of that channel. The 80% ZIKV CAM shows a higher degree of overlap with the NS4B-positive ER relative to the ERmoxGFP-, dsRNA- and NS2B-labeled ER (Figure 9A, left). Further, based on Euclidean distance analysis, the center of mass of the ZIKV CAM was closer to the NS4B center of mass (Figure 9A, right). Relative to NS2B and dsRNA, NS4B density is present throughout the CER and increased in the PER (Figure 5). Therefore, to further define the ER zone contributing to CNN discrimination of ZIKV-infected cells, we assessed overlap of the ZIKV CAM with the dsRNA-positive and dsRNA-negative CER as well as the PER, as defined previously (Figure 3). The PER, determined based on Z-height of ER segments, includes the subnuclear ER and to ensure we were assessing strictly the PER we segmented out the nuclear region using a semi-automated annotation approach (See Materials and Methods). As seen in Figure 9B, the ZIKV CAM shows increased overlap and proximity with the dsRNA-positive CER relative to either the dsRNA-negative CER or PER. The region closest to and most overlapping with the ZIKV CAM is the nuclear region, cropped from the PER channel. As the center of mass of the ZIKV CAM presents the shortest Euclidean distances to the dsRNA-positive CER and the nucleus, this locates the ZIKV CAM center of mass to the perinuclear CER region between the dense dsRNA-positive tubular matrix and the nucleus.

**Figure 9.**
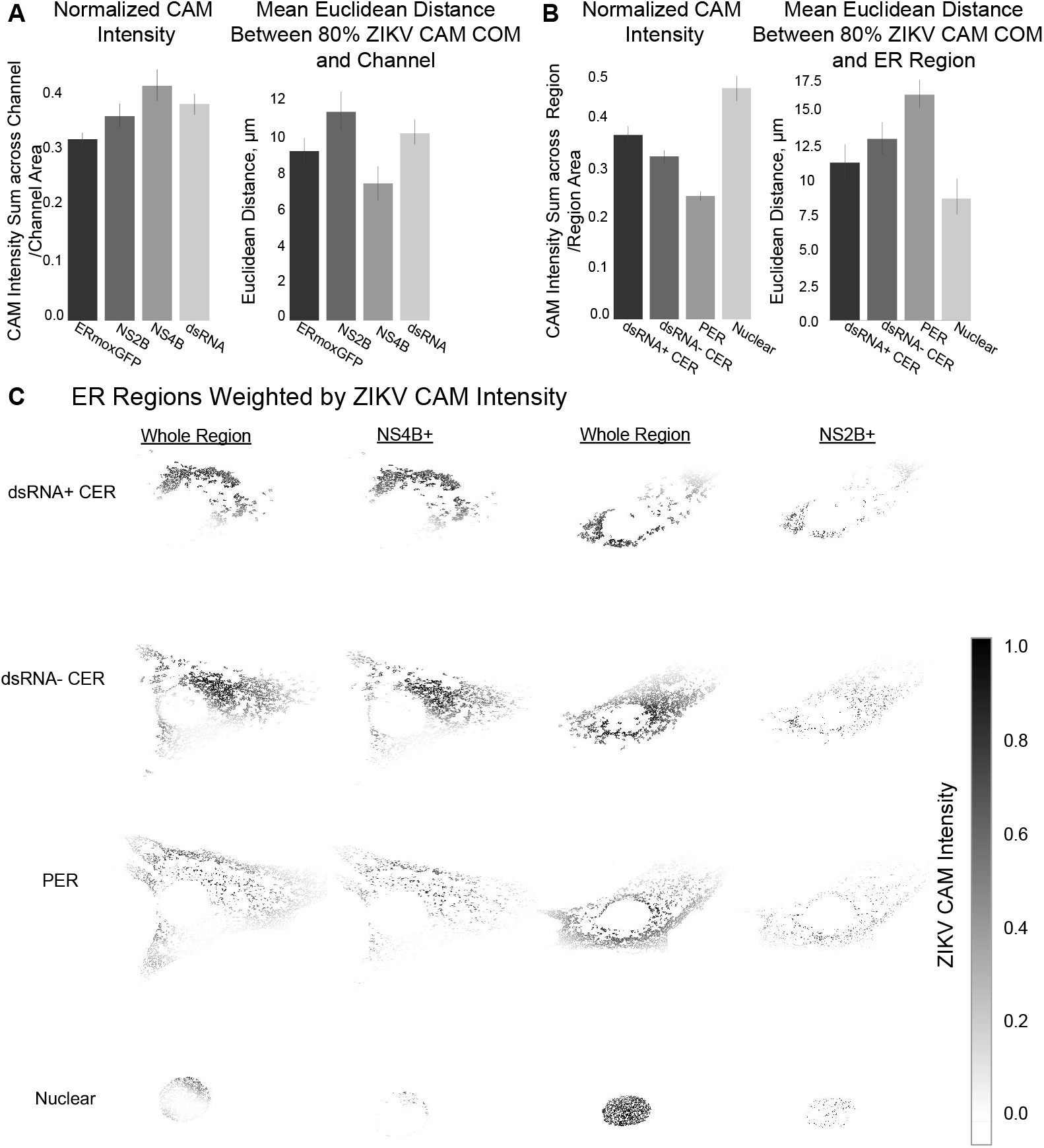
ZIKV CAM intensity relative to ER Regions. A) ZIKV CAM localization with respect to ERmoxGFP, dsRNA, NS4B and NS2B labelling. Left: Normalized CAM intensity refers to the total ZIKV CAM intensity colocalized with a given channel, normalized by the channel area. Mean Euclidean distances measured between the 80% ZIKV CAM region center of mass and all pixels of the indicated labelled protein. B) ZIKV CAM localization with respect to ER regions: dsRNA-positive CER (dsRNA+ CER), dsRNA-negative CER (dsRNA-CER), PER with nuclear region removed (PER), segmented nuclear PER region (nucleus). Same metrics are used as (B), replacing channels with ER regions. C) Representative images of NS2B and NS4B labelled ZIKV-infected cells with ER regions weighted by ZIKV CAM intensity. Darker pixels (closer to 1.0) reflect presence of ERmoxGFP and high ZIKV CAM intensity.

This can be visualized in Figure 9C, where increased grey scale density highlights the weighted overlap of the ZIKV CAM with these four ER regions. Increased ZIKV CAM overlap with the dsRNA-positive ER, the perinuclear zone of the dsRNA-negative CER as well as the nuclear zone adjacent to this region can be clearly observed. The ZIKV CAM encompasses the dsRNA, NS2B and NS4B-positive dense tubular matrix region that houses ZIKV replication factories, but also includes the adjacent perinuclear region enriched for NS4B but not NS2B. Individual patches do not encompass the totality of the CER (see Figure 7C) and the precise nature of the features that the CNN uses to discriminate between ZIKV- and mock-infected cells remains to be determined. CAM localization analysis shows that the neural network uses the same CER region that we have observed to be modified upon ZIKV infection. Deep learning therefore has the ability to identify the ER morphological changes associated with ZIKV infection.

## Discussion

ZIKV infection is characterized by re-organization of the ER to create replication factories and convoluted ER membranes involved in viral replication, whose ultrastructure has been elegantly characterized by EM (19, 25). Here, 3D STED super resolution microscopy reveals the formation of a novel, perinuclear, crescent-shaped region of dense tubular ER, or convoluted membranes, within the CER of ZIKV-infected cells. This region is enriched for dsRNA and ZIKV non-structural proteins NS2B and NS4B and therefore corresponds to the site of ZIKV replication factories and genomic replication (19, 20). 3D STED microscopy therefore shows that ZIKV infection induces CER reorganization. Importantly, based in large part on the formation of this dense CER region, deep learning image analysis is able to identify ZIKV-infected cells based solely on ER morphology. This highlights the potential utility of ER reorganization as a sensitive host cell reporter for use in screens for inhibitors of infection by ZIKV and other flaviviruses, as well as coronaviruses, such as severe acute respiratory syndrome coronavirus 2 (SARS-CoV-2), that induce extensive ER rearrangement (15, 42).

The ER is a morphologically complex organelle, containing smooth ER tubules and ribosome-studded rough ER sheets identified ultrastructurally by EM since over 60 years (43). In confocal images of cultured cells, these morphological structures correspond, respectively, to PER tubules and the dense perinuclear CER (3). Here, we approximate CER and PER regions based upon Z bounding box height of segmented 3D STED ER volumes and show that the dense ER zone enriched for dsRNA is within the CER. The CER of both ZIKV- and mock-infected U87 cells is composed of tubular networks that correspond to previously described PER tubular matrices (7). While we were unable to detect ER sheets by super-resolution analysis of cultured U87 cells, EM of brain organoids shows the transformation of ER sheets to convoluted membranes upon ZIKV infection. This suggests that organoid structures present more highly developed ER structures than cultured cells; application of 3D live cell super-resolution analysis (44) to this model of the developing fetal brain, composed of a heterogenous population of cell types, may lead to better definition of complex ER structures and their dynamic transitions in response to stress, such as viral infection. Nevertheless, the fixed cell 3D STED analysis applied here demonstrates that convoluted membranes associated with ZIKV replication derive from tubular matrix reorganization in the CER.

The ZIKV-induced CER-localized, high ER density tubular matrices are enriched for dsRNA. As dsRNA is specifically associated with viral replication, this localizes active replication factories to this CER domain that therefore corresponds to convoluted membranes (19). Functionally, the formation of tubular matrix is suggested to stabilize high curvature regions of the ER with increased membrane surface area that act as a reservoir for membrane proteins and lipids (7). ZIKV-induced dense tubular matrix may serve to provide material required for formation of replication factories and viral replication. Previously, tubular matrices were shown to shaped by both RTN and ATL proteins (Nixon-Abell et al., 2016). Consistently, depletion of RTN3.1A, ATL2 and ATL3 have been shown to reduce ER reorganization and replication of multiple flaviviruses, including ZIKV (25, 45, 46). Here, NS2B and NS4B are shown to associate with the ZIKV-induced tubular matrix and colocalize with dsRNA, localizing these proteins to replication factories. NS2B distribution is highly restricted to the ZIKV-induced tubular matrix, consistent with NS2B recruitment of NS3 to the ER and its role as a cofactor for NS3 protease function, necessary for ZIKV genomic replication and polyprotein cleavage (25, 28, 47). In contrast, NS4B has been shown to be a multifunctional, membrane protein involved in many processes of ZIKV infection and pathogenesis, including promoting ER membrane proliferation, curvature of ER membranes to produce replication factories and disruption of ER-mitochondria contacts allowing the virus to successfully subvert the host innate immune response (26, 48, 49). Demonstration here that NS4B presents a broad distribution throughout the ER is consistent with such a role and implicates NS4B in the ER reorganization that leads to tubular matrix formation. A complete understanding of the spatial and temporal role of NS and ER shaping proteins in ZIKV-induced tubular matrix formation remains to be determined.

We show that deep convolutional neural networks can accurately classify ZIKV- and mock-infected cells based on ER labeling. Through the use of CAMs, we focus on regions which are more likely to be used by the network to infer a class label. Importantly, our analysis shows that the VGG16 neural network uses dense ERmoxGFP in the CER as an indicator of ZIKV infection. This demonstrates that the neural network is detecting the ZIKV-induced reorganization of the CER that leads to the formation of the tubular matrix associated with ZIKV replication. Accuracy obtained for VGG16 classification of ZIKV-infected cells from 3D STED image stacks is comparable to prior classification using VGG16. VGG16 achieved 79.2% accuracy when classifying disease (e.g., melanoma) versus benign skin lesions from the ISIC 2016 challenge dataset of dermoscopy images (37, 50). Analysis of dystrophic MRI data classified Duchenne and congenital muscular dystrophies with an accuracy of 84.1% using VGG16, which was boosted by 7.6% when implementing their improved CAM method into a ResNet50 base model (51). For an entirely different task and dataset, VGG16 achieved highest accuracy (92.6%) compared to ResNet50 (79.4%), when trained to classify histopathological breast cancer images (as benign or malignant breast tissue) from the publicly available BreakHis Dataset (38, 52). Here, our analysis of a novel 3D STED data set introduces a baseline for classification performance for 3D super-resolution microscopy data sets that will be of service to other researchers applying deep learning approaches to super-resolution microscopy.

The interpretability of artificial intelligence is an evolving field and we believe that interpretable methods, such as Grad-CAM (53), are important tools for the understanding of deep neural networks applied to exploratory data sets. This approach has now allowed us to identify features of discriminatory regions, and has not, to our knowledge, been applied to subcellular morphology, nor to 3D super resolution images of the ER. Comparison of downsampled low-resolution images from 2D sections of cells presenting ER reorganization into a denser CER tubular matrix was sufficient to train the neural network to identify ZIKV-infected cells. This highlights the sensitivity of deep learning image analysis and augurs well for future identification not only of tubular matrix but also of the peripheral tubules, 3-way junctions, sheets and organelle contacts that comprise the ER (54). However, defining the morphological characteristics associated with deep learning based decision making from isolated 2D patches of ER remains a challenge. Future application of similar deep learning approaches to super-resolution 3D analysis of both fixed and live cells may lead to novel understanding of the dynamic, local changes to ER morphology associated with the diverse functions of this cellular organelle. To the best of our knowledge, interpreting the network’s decisions is a novel approach to study ER morphology using deep learning.

Screening of siRNA or small molecule libraries to identify inhibitors of viral infection is a valid and urgent research endeavour, given the current SARS-CoV-2 pandemic. Such high-throughput screens require robust, sensitive assays to identify potential inhibitors for subsequent secondary screens and drug development. The intent of this study is not the use of deep learning to identify virus-infected cells (55) but rather the use of deep learning to identify and localize the morphological features of the ER. This includes but is not limited to the central ER tubular matrices induced by ZIKV infection and therefore, in combination with 3D super-resolution microscopy, has tremendous potential to identify distinct, local morphological features of the ER and other cellular organelles. Nevertheless, application of a deep learning framework to host-cell ER labeling correctly identified ZIKV-infected cells from 2D sections of downsampled STED super-resolution images; this indicates that VGG16 and other deep CNN architectures are capable of finding discriminatory features in lower resolution images for correct identification of infected cells. Importantly, our deep learning approach is translatable to ER reorganization induced by other pathogenic human enveloped viruses. As for flaviviruses (i.e. ZIKV), the ER is the key replication site for coronaviruses, likely including SARS-CoV-2 responsible for COVID-19, whose infection is also associated with ER reorganization to form convoluted membranes (15, 42, 56). Deep learning analysis of host cell reorganization of the ER may therefore represent a sensitive detection approach to identify virus infected cells.

## Methods and Materials

### Antibodies, plasmids, chemicals and viruses

ERmoxGFP was a gift from Dr. Erik Snapp (Albert Einstein College of ERmoxGFP was a gift from Dr. Erik Snapp (Albert Einstein College of Medicine, presently at Howard Hughes Medical Institute, Janelia Research Campus, Virginia) (Addgene plasmid #68072), and Sec61β-GFP from Dr. Gia Voeltz (University of Colorado, Boulder). Mouse anti-dsRNA antibody (Cat#: J2-1904) was purchased from Scicons English and Scientific Consulting, goat anti-rabbit Alexa Fluor 532 from ThermoFisher Scientific (Cat#: A-11009), goat anti-mouse Alexa Fluor 568 from ThermoFisher Scientific (Cat#: A-11031), and goat serum from Thermo Fisher Scientific (Cat#: 16210-064). Rabbit anti-NS2B (Cat#: GTX133308) and NS4B (Cat#: GTX133321) were kindly provided by Genetex (Cat#: GTX133321). 16% paraformaldehyde (Cat#: 15710) and 25% glutaraldehyde (Cat#: 16220) were from Electron Microscopy Sciences, USA. All other chemicals were obtained from Sigma. ZIKV strain (PRVABC59) was obtained from ATCC (Cat#: VR-1843).

### Cells and organoids

U87 MG cells from ATCC (Cat#: HTB-14) were grown at 37°C with 5% CO2 in Gibco Minimum Essential Media (MEM) (Cat#: 11095080) containing 10% FBS, 1% MEM non-essential amino acids (Cat#: 11140050), and 1% sodium pyruvate (Cat#: 11360070) (ThermoFisher Scientific) unless otherwise stated. Plasmids were transfected in U87 cells with Effectene (Qiagen, Germany) according to the manufacturer’s protocols. Following 24 hours of plasmid transfection cells were incubated in MEM with 800 mg/mL of Geneticin (Cat#: 10131027) to select for transfected cells. Media was changed every 24 hours for a week and then every 2 days. Once colonies grew to confluency the resultant GFP positive cells were sorted by FACS. Plasmid stable lines were maintained in MEM with 400 mg/mL of Geneticin where media was changed every two days. Human pluripotent stem cell (hPSC)-derived cerebral organoids were grown at 37°C with 5% CO2 in STEMdiff Cerebral Organoid Kit cell culture media from STEMCELL Technologies Inc. (Cat#: 08570).

### Immunofluorescence labeling

Cells grown on #1.5H coverslips (Paul Marienfeld GmbH & Co, Germany) for 24 hours were washed once with phosphate buffered saline (PBS) and incubated with ZIKV, diluted in MEM, at a MOI of 1 for 48 hours. Then cells were: 1) fixed with 3% paraformaldehyde with 0.2% glutaraldehyde at room temperature for 15 minutes and washed with PBS-CM (PBS supplemented with 1 mM CaCl2 and 10 mM MgCl2) (all PBS-CM washes consist of two quick washes and then two 5 minute washes unless otherwise specified); 2) permeabilized with 0.2% Triton X-100 for 5 minutes followed by PBS-CM washes; 3) quenched of glutaraldehyde aldehydes with 1 mg/mL of NaBH4 for 10 minutes and washed with PBS-CM; 4) blocked with 10% Goat serum (ThermoFisher Scientific) and 1% bovine serum albumin (Sigma) in PBS-CM for 1 hour; 5) incubated with primary antibodies in the same blocking solution as described above for 1 hour at room temperature (RT) then washed quickly with PBC-CM once followed by three 5 minute washes with Antibody Wash Buffer (20x SSC, 0.05% Triton-X100 in Milli-Q H2O); 6) incubated with secondary antibodies in Antibody Buffer (1% BSA, 2% goat serum, 0.05% Triton-X100, 20X sodium/sodium citrate buffer in Milli-Q H2O) for 1 hour at RT then washed quickly with PBS-CM once followed by six 10 minute washes with Antibody Wash Buffer on a rocker; 7) rinsed with Milli-Q H2O and mounted with ProLong Diamond (ThermoFisher Scientific) and cured for 24-48 hours at RT (6).

### 3D STED microscopy

3D STED imaging was performed with the 100X/1.4 Oil HC PL APO CS2 objective of a Leica TCS SP8 3X STED microscope (Leica, Germany) using white light laser excitation with HyD detectors, and Leica Application Suite X (LAS X) software. Sample acquisition was done at a scan speed of 600Hz with a line average of 6. Time-gated fluorescence detection was used to further improve lateral resolution. GFP was excited at 488 nm and depleted using the 592 nm depletion laser. Alexa Fluor 532 was excited at 528 nm, and Alexa Fluor 568 was excited at 577 nm. Both were depleted using the 660 nm depletion laser. To avoid crosstalk, image stacks for each channel were acquired sequentially, in the order of Alexa Fluor 568, Alexa Fluor 532 and then GFP. A step size of 210 nm was utilized for 3D STED. Images were then deconvoluted using Huygens Professional software (Scientific Volume Imaging). 3D STED ER segmentation and ER region mask generation were done using Imaris x64 9.2.1. dsRNA associated ER and NS4B/NS2B associated ER regions were generated by partial overlap analysis after surface generation. All data produced from Imaris were parsed and analyzed with a custom made Jupyter notebook. Two-tailed unpaired Student’s t-tests or ANOVA with post-hoc Tukey HSD were done using Graph Pad Prism 6.0.

### Transmission electron microscopy

63-day old cerebral organoids were infected with the PRVABC59 ZIKV strain, MOI of 1, for 48 hours. Organoids were then processed by replacing media with the primary fixative [0.1 M sodium cacody-late, 1.5% paraformaldehyde, 1.5% glutaraldehyde, pH 7.3, room temperature] (57) and allowed to sit for 1 hour before placing them at 4°C overnight. The organoids were washed three times (10 minutes each wash) with buffer (0.1M sodium cacodylate) and then post-fixed for 1 hour on ice in 0.1M osmium tetroxide in 0.1M sodium cacodylate (pH7.3). They were then washed three times with distilled water (10 min each wash, room temperature) and stained en bloc with 1% aqueous uranyl acetate for 1 hour at room temperature. They were washed again three times with distilled water and then dehydrated through an ascending concentration series of ethyl alcohol (30%, 50%, 70%, 95%, 100%), passed through two changes of propylene oxide, and then infiltrated overnight in a 1:1 solution of propylene oxide:EMBED 812 resin. The organoids were then passed through two changes of 100% EMBED resin, embedded and the resin polymerized at 60°C for 48 hours. Thin sections were cut using a diamond knife and sections were collected on copper grids. The sections were then stained with uranyl acetate and lead citrate, and images collected using a FEI tecnai G2 Spirit transmission electron microscope operated at 120 kV.

### Image data processing for CNN analysis

A custom pipeline was created using MATLAB R2019a (The Mathworks, Inc., Natick, MA) and Python 3 (Python Software Foundation, Scotts Valley, CA) scripts to convert data stored in Leica proprietary format (.lif) to an accessible and lightweight format, which could easily feed as input to state of the art CNNs. Once an imaging session is complete, the biologist exports Leica STED data without need for further processing or file conversion. The pipeline takes Leica .lif files of 3D STED volumes of multiple cells as input and outputs a single NumPy array per cell. The pipeline relies on open-source Matlab package Bio-Formats, modified to handle our needs. This pipeline has been validated through manually inspection of cells with Imaris image rendering software.

### Deep learning

The Keras open-source neural network library was used to modify and train neural networks using Tensorflow backend. VGG16 was used as a base model to predict infected from non-infected 2D frames (removing the original fully-connected output layers). This architecture and pretrained weights (learned using the ImageNet dataset) are easily accessible using Keras. The pretrained weights can be loaded when instantiating the base model. On top of the VGG16 max pooling layer, we add (in order): a global average pooling layer, a dropout layer (with a dropout rate of 0.5), dense layer (using ReLu activation, output dimension 1024), a second dropout layer (0.5 dropout rate) followed by a final dense layer (with softmax activation) for making predictions. These additional layers helped prevent overfitting of the CNN to the training data.

56 individual CNNs (with identical architecture as outlined above) were trained using a softmax loss function and RMSprop optimizer. Each model was then used to predict the class labels (i,e. assign either ‘Mock’ or ‘ZIKV’ to each 2D frame) of a single test cell. The CNNs were fine-tuned for a total of 24 epochs, after which only the best weights (resulting in highest classification performance on the validation set) were then reloaded during test time. CAMs were generated following the procedure outlined by (39), where global average pooling is applied to the output of the final convolutional layer of VGG16 followed by a weighted sum, according to the specified class (e.g., ZIKV). Models were trained on 2 Nvidia GPUs (GeForce GTX TITAN X, each with 32 GB of RAM).

ROIs generated through the deep learning framework were analyzed using python programming. Statistical analysis across channels and ER regions relied on several open-source python packages. SciPy’s statistical functions module, was used to perform Kruskal-Wallis non-parametric tests, statsmodules python module was used for One-way ANOVA, and scikit-posthocs python package was used to perform Dunn’s post hoc test.

### Code availability

All code related to data processing and deep learning and was written using open-source packages.

## Acknowledgments

This study was supported by grants to IRN from the Canadian Institutes for Health Research (CIHR; PJT-148698) and Natural Sciences and Engineering Research Council of Canada (NSERC; RGPIN-2019-05179), to GH from NSERC (RGPIN-2015-06795, RGPIN-2020-06752), and to FJ from CIHR (22R76798). RL is the recipient of a CIHR Frederick Banting and Charles Best Canada Graduate Scholarship-Master’s award (CGS M). STED imaging and analysis was performed in the LSI IMAGING facility of the Life Sciences Institute of the University of British Columbia using infrastructure funded by the Canadian Foundation of Innovation and BC Knowledge Development Fund as well as a Strategic Investment Fund (Faculty of Medicine, University of British Columbia). We further thank Dr. Alex Ball, Jr., M.D. Senior Scientist at Genetex, for supplying the rabbit anti-NS4B and anti-NS2B antibodies and Leon Chew, STEMCELL Technologies Inc, for supplying the cerebral organoids used in this study. We thank Compute Canada for providing computational resources and NVIDIA Corporation for donating the GeForce GTX TITAN X GPUs used in this research.

## Supplemental Material

Supplemental Figure I: Network Performance Analysis

**Figure 10.**
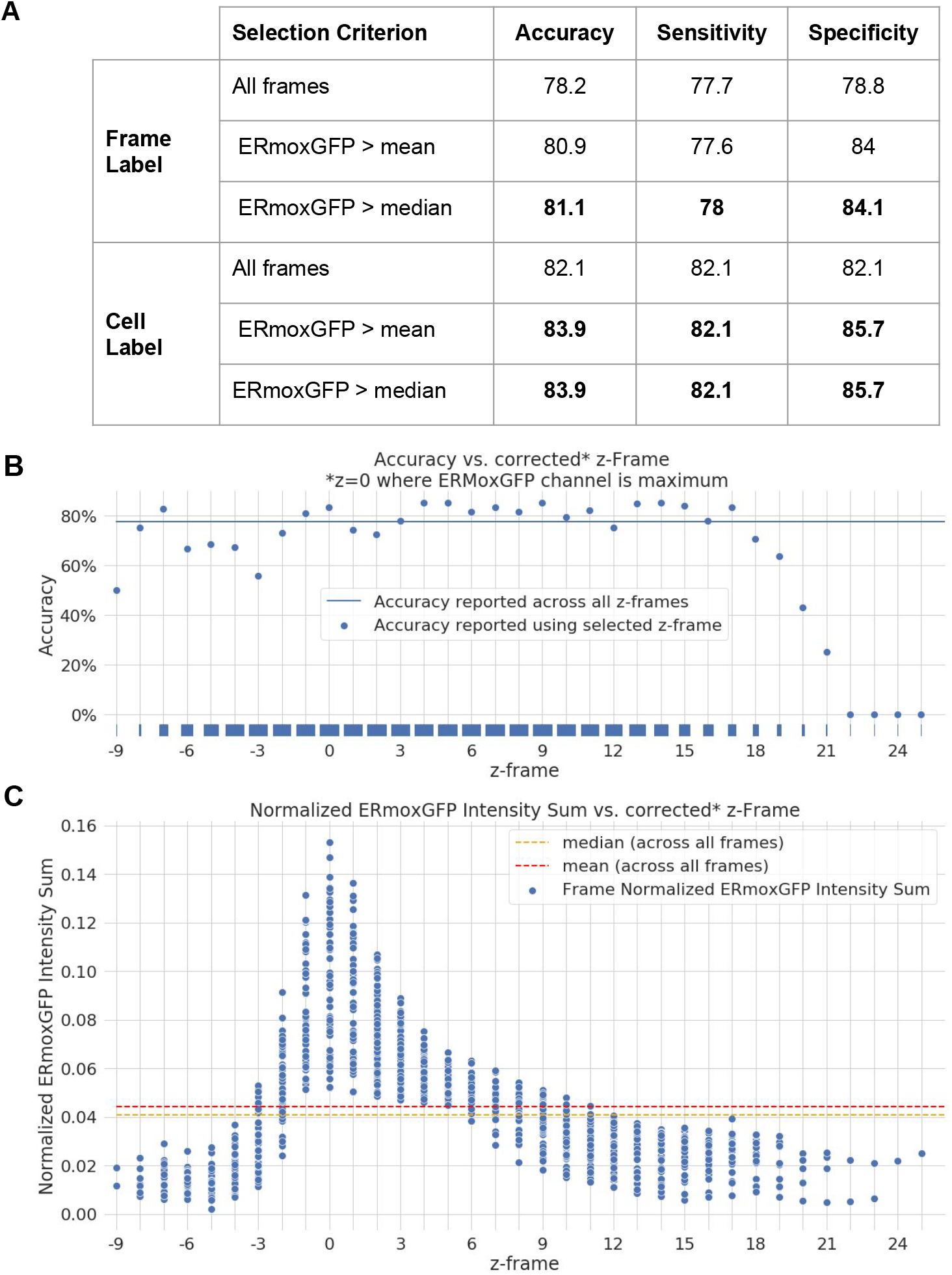
Network Performance Analysis A) Performance metrics are reported across predictions of frame labels and cell labels, where cell labels correspond to the majority label of predicted frame labels for each cell. Results are reported using a given selection criteria: using all frames (rows 1, 4), using only frames with normalized ERmoxGFP signal greater than mean normalized ERmoxGFP signal (rows 2, 5) or greater than median normalized ERmoxGFP signal (rows 3, 6). Mean and median thresholds are computed on a cell basis. The rug plot (above x-axis) visualises distribution of z-frames. B) Accuracy reported across corrected z-frames, z=0 is where normalized ERmoxGFP Intensity Sum is maximal. C) Normalized ERmoxGFP Intensity sum plotted against corrected z-frame. Dashed lines indicate the median (orange) and mean (red) Normalized ERmoxGFP Intensity sum computed across all frames.

